# Functional magnetic resonance imaging of the lumbosacral cord during a lower extremity motor task

**DOI:** 10.1101/2024.01.31.577917

**Authors:** Christian W. Kündig, Jürgen Finsterbusch, Patrick Freund, Gergely David

**Author notes:** Corresponding author. Gergely David, Spinal Cord Injury Center Balgrist University Hospital Forchstrasse 340, 8008 Zurich, Switzerland.

## Abstract

Blood-oxygen-level dependent (BOLD) functional magnetic resonance imaging (fMRI) can be used to map neuronal function in the cervical cord, yet conclusive evidence supporting its applicability in the lumbosacral cord is still lacking. This study aimed to (i) demonstrate the feasibility of BOLD fMRI in mapping neuronal activation in the lumbosacral cord during a unilateral lower extremity motor task and (ii) investigate the impact of echo time (TE) on the BOLD effect size. Twelve healthy volunteers underwent BOLD fMRI using four reduced-field-of-view single-shot gradient-echo echo planar imaging sequences, all with the same geometry but different TE values ranging from 20 to 42 ms. Each sequence was employed to acquire a single 6-minute rest run and two 10-minute task runs, which included alternating 15-second blocks of rest and unilateral ankle dorsi- and plantar flexion. We detected lateralized task-related neuronal activation at neurological levels S4 to L1, centered at the ipsilateral (right) ventral spinal cord but also extending into the ipsilateral dorsal spinal cord. This pattern of activation is consistent with our current understanding of spinal cord organization, wherein lower motor neurons are located in the ventral gray matter horn, while sensory neurons of the proprioceptive pathway, activated during the movement, are located in the dorsal horns. At the subject level, BOLD activation showed considerable variability but was lateralized in all participants. The highest BOLD effect size within the ipsilateral ventral spinal cord was observed at TE=42 ms. Sequences with a shorter TE (20 and 28 ms) also detected activation in the medioventral part of the spinal cord, likely representing a large vein effect. In summary, our results demonstrate the feasibility of detecting neuronal activation in the lumbosacral cord induced by voluntary lower limb movements. BOLD fMRI in the lumbosacral cord has significant implications for assessing motor function and its alterations in disease or after spinal cord injury.

## 1. Introduction

Functional MRI (fMRI) has been increasingly applied in the spinal cord to non-invasively map spinal cord neuronal function (Cohen-Adad, 2017; Haynes et al., 2023; Landelle et al., 2021; Powers et al., 2018). Spinal cord fMRI conducted in healthy volunteers have investigated neuronal activation induced by a wide range of tasks and stimuli, including upper extremity motor task (Govers et al., 2007; Kinany et al., 2019; Stroman et al., 1999; Weber et al., 2016b), thermal stimulation (Bosma & Stroman, 2015; Rempe et al., 2015; Weber et al., 2016a, 2018), tactile stimulation (Brooks et al., 2012; Kornelsen et al., 2013; Summers et al., 2010; Weber et al., 2020), noxious stimuli (Brooks et al., 2012; Summers et al., 2010; Tinnermann et al., 2021), and sexual arousal (Alexander et al., 2016; Kozyrev et al., 2012). Importantly, studies have reproduced the expected lateralization of spinal cord neuronal activation in response to ipsilateral tasks (Kinany et al., 2019; Weber et al., 2016b) or stimulation (Brooks et al., 2012; Weber et al., 2020), along with the expected rostrocaudal distribution (Kinany et al., 2019). Spinal cord fMRI has also been applied in the absence of explicit tasks (i.e., resting state) to uncover the intrinsic functional networks within the spinal cord (Barry et al., 2014, 2016; Combes et al., 2023; Kinany et al., 2020; Vahdat et al., 2020; Weber et al., 2018).

The majority of investigations thus far have focused on the cervical cord due to (i) the relatively high signal-to-noise ratio (SNR) facilitated by optimal coverage provided by the head and neck coils, (ii) the comparatively large cross-section of the cervical spinal cord, (iii) the relatively straight shape of the upper cervical cord, and (iv) the ease of minimizing task-related motion artifacts in the spinal cord during upper extremity motor tasks. In contrast, the lumbosacral cord, a region crucial for functions such as locomotion, reflexes, as well as sexual, bladder, and bowel control (Fowler et al., 2008; Krassioukov & Elliott, 2017; Sharrard, 1964), has received limited attention. This discrepancy can be attributed primarily to the smaller size of the lumbosacral cord and the technical challenges it presents, including the lower SNR associated with the spine coils and difficulties in shimming due to the proximity of large vertebrae and the lungs. Indeed, out of the 44 papers identified in a literature review (Landelle et al., 2021), only one focused on the lumbosacral cord. In that study, motor and sensory tasks were performed on 6 subjects using a 1.5T MRI scanner, utilizing the signal enhancement by extra-vascular water proton (SEEP) contrast mechanism (Kornelsen & Stroman, 2004). In a recent study, blood-oxygen-level-dependent (BOLD) fMRI was employed to map neuronal activation in the lumbosacral cord during passive movements and muscle tendon vibration of the lower limbs in three patients with spinal cord injury (Rowald et al., 2022). Although yielding promising results, definitive evidence supporting the applicability of BOLD fMRI in the lumbosacral cord is still lacking.

A significant challenge in spinal cord fMRI is the lack of standardized imaging protocols, which limits the comparability of findings. While generic acquisition protocols and guidelines exist for quantitative MRI of the spinal cord (Cohen-Adad et al., 2021a, 2021b), a comparable framework for spinal cord fMRI has yet to be established. Achieving this standardization necessitates a comprehensive understanding of how sequence parameters impact the fMRI results. In this context, Kinany and colleagues conducted a comparative analysis of frequently used fMRI sequences, evaluating their efficacy in detecting task-related activation and resting-state functional networks (Kinany et al., 2022). However, their investigation focused on the cervical cord, which might not generalize to the lumbosacral cord. Therefore, there is a need to provide recommendations for optimal BOLD fMRI in the lumbosacral cord.

The goal of our study is twofold. First, we aimed to demonstrate the feasibility of lumbosacral BOLD fMRI by using a T2*-weighted single-shot gradient-echo echo planar imaging (GE-EPI) sequence during a unilateral lower extremity (ankle) motor task. Second, we investigated the influence of echo time (TE), an important determinant of BOLD sensitivity (Menon et al., 1993; Triantafyllou et al., 2011), on the level and extent of BOLD activation within the lumbosacral cord.

## 2. Methods

### 2.1 Subjects

Twelve healthy subjects (4 females, 8 males, age (mean ± standard deviation (SD)): 28.4 ± 3.4 years) participated in the study. All subjects were right-footed. The study received approval from the Cantonal Ethics Committee of Zürich (BASEC ID: 2022-00558), and written informed consent was obtained from all participants.

### 2.2 Image acquisition

#### 2.2.1 Hardware and subject positioning

The scanning was performed using a 3T whole-body MR system (Magnetom Prisma, Siemens Healthineers) with a body transmit coil for excitation and a 32-channel spine matrix coil for reception. A foam wedge was placed under the legs to reduce the natural lordotic curve and maximize contact with the spine matrix coil. To minimize any movement of the lower spine due to the ankle motion task or involuntary motion, we (i) placed a vacuum cushion under the legs, (ii) applied body straps around the knees and hips, and (iii) instructed the participants to avoid heavy breathing. The ankle positioning allowed participants to perform the full range of ankle dorsiflexion and extension without touching the foam wedge.

#### 2.2.2 Anatomical scans

To facilitate the slice prescription for the subsequent axial images, a sagittal T2-weighted turbo spin echo sequence with 15 slices of 4 mm thickness (10% slice gap) was acquired, encompassing the lumbosacral enlargement (LSE) and the conus medullaris (Fig. 1A,C). Additional sequence parameters were: in-plane field of view of 330×330 mm^2^, in-plane resolution of 0.7×0.7 mm^2^, repetition time (TR) of 3000 ms, echo time (TE) of 89 ms, echo spacing of 8.94 ms, turbo factor of 16, flip angle (FA) of 151°, phase oversampling of 75%, parallel imaging with GRAPPA (acceleration factor of 2), bandwidth of 272 Hz/pixel, and acquisition time of 00:59 min.

**Fig. 1.**
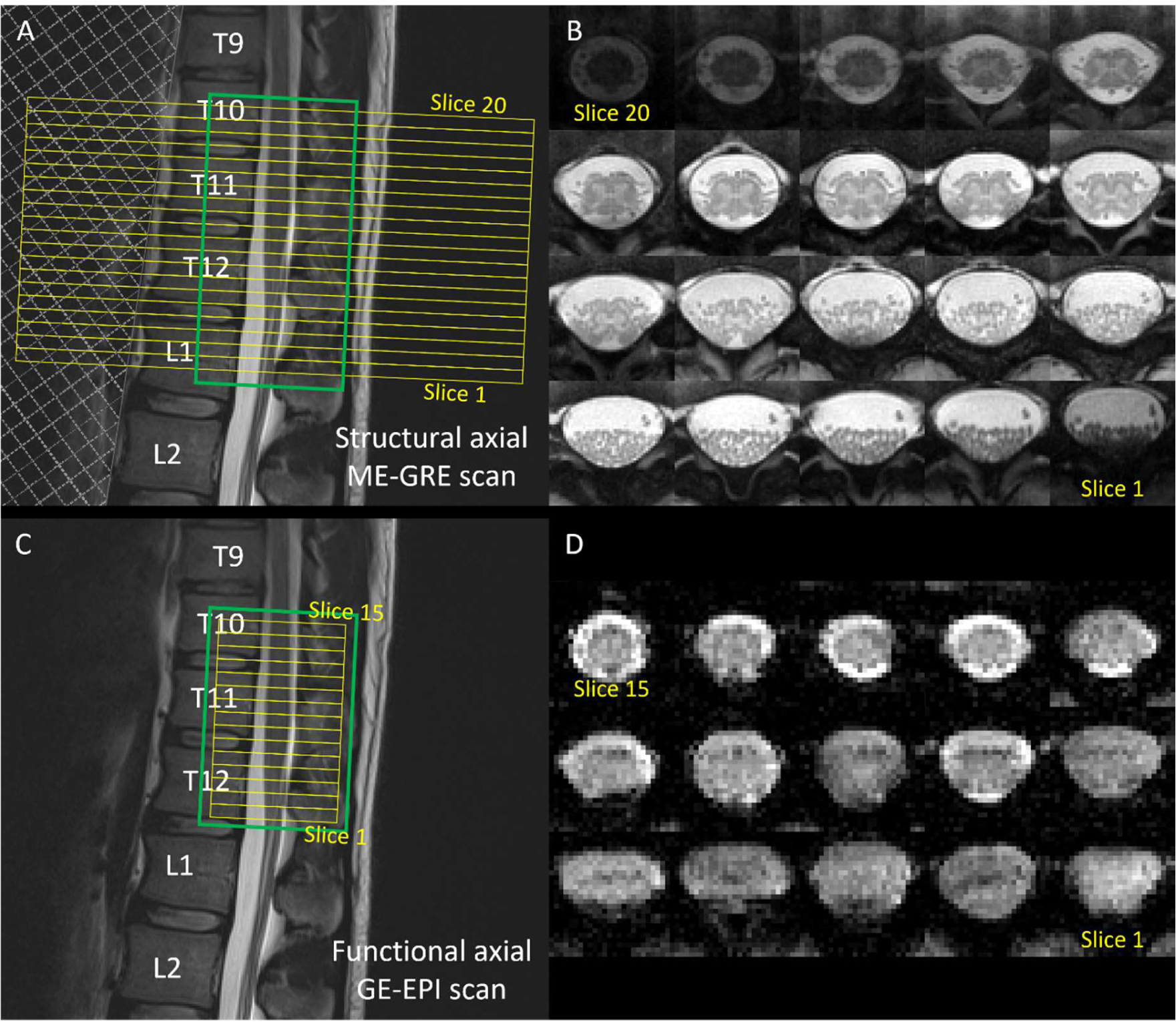
Slice prescription, based on a sagittal T2-weighted turbo spin echo scan of the lower spine, for the structural 3D spoiled multi-echo gradient-echo (ME-GRE, Siemens FLASH) sequence (A) and the functional reduced-field-of-view single-shot gradient-echo echo planar imaging (GE-EPI) sequences (C). The slice stacks (20 slices for ME-GRE, 15 slices for GE-EPI) and shimming volumes are indicated by yellow and green boxes, respectively. The field of view (FOV) for the GE-EPI scan was set such that its 6^th^ most rostral slice (slice #10) aligned with the maximum width of the spinal cord in the lumbosacral enlargement as observed in the sagittal T2-weighted image. The FOV for the ME-GRE scan was positioned to align the top edge with that of the GE-EPI scan (i.e., slice #20 in ME-GRE corresponds to slice #15 in GE-EPI), which ensured coverage of the lumbosacral enlargement and the conus medullaris. For the ME-GRE sequence (A), a saturation band, indicated by a shaded area, was placed anterior to the spinal column to suppress potential artifacts originating from the abdomen. Note that the OVS20 version of the GE-EPI sequence (Table 1) also included two saturation bands, placed anterior and posterior to the FOV. The corresponding axial slices are shown for the ME-GRE (B) and GE-EPI sequences (here, the iFOV35 version, see Table 1) (D), displayed in rostral (top left) to caudal (bottom right) direction.

**Table 1.** Sequence parameters of the single-shot gradient-echo echo planar imaging sequences used for functional MRI.

| Sequence | OVS20 | iFOV28 | iFOV35 | iFOV42 |
| --- | --- | --- | --- | --- |
| *Varying parameters* |  |  |  |  |
| Type of reduced FOV | OVS | iFOV | iFOV | iFOV |
| Partial Fourier factor | 6/8 | 6/8 | 7/8 | no |
| Echo time (ms) | 20 | 28 | 35 | 42 |
| Repetition time (ms) | 1460 | 1190 | 1290 | 1400 |
| Flip angle (°) | 77 | 72 | 74 | 76 |
| Number of volumes (task) | 411 | 505 | 466 | 429 |
| Number of volumes (rest) | 247 | 303 | 280 | 258 |
| *Identical parameters* |  |  |  |  |
| Number of slices |  | 15 |  |  |
| Slice thickness (mm) |  | 5.0 |  |  |
| Slice acquisition |  | interleaved |  |  |
| FOV (mm<sup>2</sup>) read x phase (L-R x A-P) |  | 144 x 48 |  |  |
| Resolution (mm<sup>2</sup>) |  | 1.0 x 1.0 |  |  |
| Echo spacing (ms) |  | 0.93 |  |  |
| PAT |  | no |  |  |
| Bandwidth (Hz/pixel) |  | 1240 |  |  |
| Acquisition time (task) |  | 10:05 – 10:07 min |  |  |
| Acquisition time (rest) |  | 06:05 - 06:07 min |  |  |

A-P, anterior-posterior direction; FOV, field of view; iFOV, inner field-of-view acquisition; L-R, left-right direction; OVS, outer volume suppression; PAT, parallel acquisition technique.

A high-resolution axial scan was acquired using a 3D spoiled multi-echo gradient-echo (ME-GRE) sequence with 20 axial-oblique slices of 5.0 mm thickness (no gap) to serve as an anatomical reference (Fig. 1A,B) for the functional scans (Fig. 1C,D). Further acquisition parameters were: in-plane FOV of 192×192 mm^2^, in-plane resolution of 0.5×0.5 mm^2^, TR of 38 ms, first TE of 6.85 ms, echo spacing of 4 ms, echo train length of 5 with bipolar readout, FA of 8°, phase encoding along the anterior-posterior direction, parallel imaging with GRAPPA (acceleration factor of 2), water excitation to avoid fat signal, flow compensation, bandwidth of 260 Hz/pixel, 6 measurements, and acquisition time of 13:28 min.

#### 2.2.3 Functional scans

Functional scans comprised four different T2*-weighted reduced-FOV single-shot GE-EPI sequences, acquired with identical geometry. The 15 axial-oblique slices of 5.0 mm thickness were acquired in an ascending interleaved order without gap. The EPI sequences had differences in TE, TR, FA, partial Fourier factor (pF), and the way reduced FOV was achieved (outer volume suppression (OVS) or inner field-of-view excitation (Finsterbusch, 2013) (refer to Table 1 for sequence parameters).

Three inner-FOV GE-EPI sequences were acquired using Siemens’ ZOOMit implementation with varying TE (hereafter: iFOV28, iFOV35, and iFOV42, where the number at the end indicates the TE in ms). Shorter TE was achieved by applying pF in the phase-encoding direction, where TE was always set to the minimum (iFOV28: pF of 6/8; iFOV35: pF of 7/8; iFOV42: no pF). In addition, a single OVS EPI sequence (hereafter: OVS20) was acquired by placing two saturation bands, each with a thickness of 130 mm, anterior and posterior to the FOV, which allowed for an even shorter TE of 20 ms in combination with pF of 6/8. For each sequence, TR was set to the minimum and FA was set to the Ernst angle, using the given TR and a T1 estimate of 994 ms within the cervical gray matter at 3T (Smith et al., 2008). The only exception was OVS20, where the TR was increased from the minimum of 1130 ms to 1460 ms to comply with the specific absorption rate limit. All other sequence parameters were identical between the four sequences, including the in-plane FOV of 144×48 mm^2^, in-plane resolution of 1.0×1.0 mm^2^, phase encoding along the anterior-posterior direction, phase oversampling of 25%, and bandwidth of 1240 Hz/pixel.

### 2.3 Experimental paradigm and study design

Fig. 2 illustrates the experimental paradigm and study design. The four GE-EPI sequences were acquired in a counterbalanced order. The anatomical scan was acquired between the second and the third GE-EPI sequence. For each GE-EPI sequence, we acquired one resting-state run followed by two task runs. The task paradigm consisted of alternating blocks of rest and right-sided ankle dorsiflexion/plantar flexion movements. The right side was chosen as it was the dominant side in all participants. Each block was 15 s long, and there were 40 blocks (20 motion and 20 rest blocks) in a task run, starting with a rest block.

**Fig. 2.**
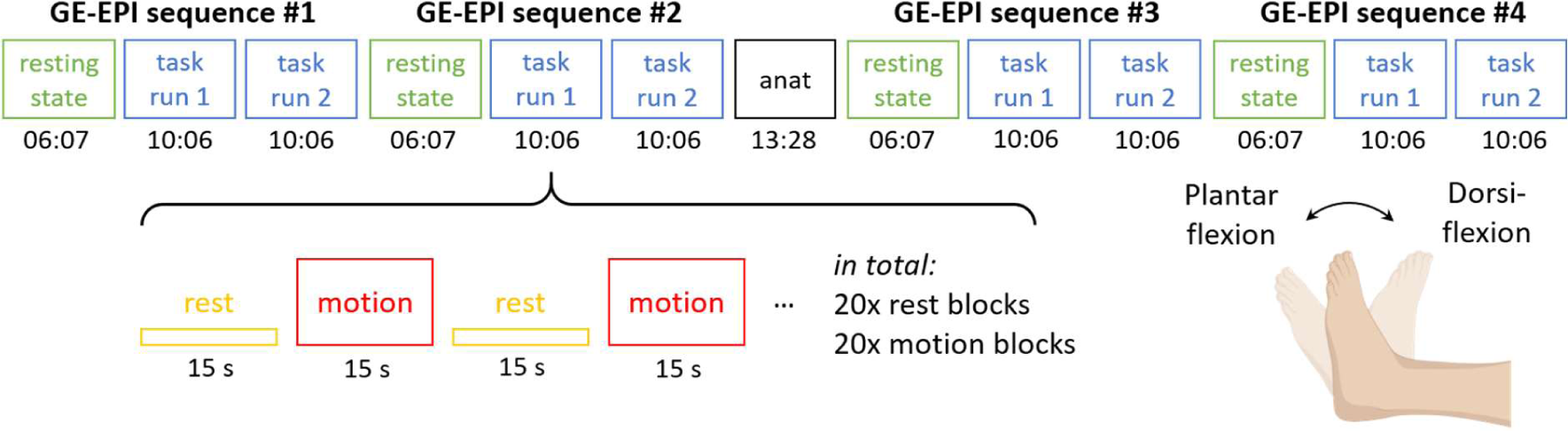
The lower extremity motor task paradigm (unilateral ankle dorsiflexion and plantar flexion). Part of the figure was created with BioRender.com.

During the task runs, participants observed the screen for instructions via a mounted mirror. A blank screen indicated rest (i.e., no ankle movement), while a flashing circle indicated ankle movement. Participants were instructed to perform the full range of dorsiflexion and plantar flexion at a constant rhythm according to the frequency of the flashing (1 Hz), meaning that participants transitioned from dorsiflexion (i.e., ankle in the highest position) to plantar flexion (i.e., ankle in the lowest position) in 1 s, and vice versa. During the rest runs, participants were instructed to lay still with their eyes open.

Note that the identical acquisition times but different TR resulted in different number of volumes for both task fMRI (OVS20: 411; iFOV28: 505; iFOV35: 466; iFOV42: 429 volumes) and resting-state fMRI (OVS20: 247; iFOV28: 303; iFOV35: 280; iFOV42: 258 volumes) (Table 1). Three dummy volumes were acquired at the start of each GE-EPI sequence, which were discarded by the scanner and did not trigger the task paradigm.

### 2.4 Processing of the anatomical images

The ME-GRE sequence yielded 30 images corresponding to five echoes in each of the six repetitions. In each repetition, we averaged all echoes into a single image (“combined echo”) by root-mean-square summation, followed by averaging across the six measurements using serial longitudinal registration (part of SPM12) to account for motion between measurements. To match the FOV of the cropped EPI volumes, the resulting image was cropped in-plane around the center of the image to a matrix size of 97×97. The cropped image was used for further analyses and is referred to as the ME-GRE image throughout the paper. The ME-GRE image was segmented manually for spinal cord using the sub-voxel segmentation tools in JIM (version 7.0, Xinapse Systems Ltd, UK). Sub-voxel segmentations were binarized with an inclusion threshold of 50%.

### 2.5 Processing of the functional images

#### 2.5.1 Motion correction

The EPI volumes were cropped in-plane around the spinal cord from a matrix size of 144×48 to 48×48 to reduce computing time and storage space. The EPI volumes were motion corrected in a two-step process. First, we ran a volume-wise motion correction with rigid-body (6 degrees of freedom) transformation using MCFLIRT, applied with least squares cost function and spline final interpolation (Jenkinson et al., 2002) to register each volume to the mean EPI volume. The spinal cord centerline was generated on the mean motion-corrected EPI volume using the *sct_get_centerline* function in Spinal Cord Toolbox (SCT) (De Leener et al., 2017). Then, we fed the output into a regularized slice-wise motion correction algorithm (*sct_fmri_moco*), implemented in SCT and applied with two degrees of freedom (translation along x and y), first-order polynomial regularization function (along the slice-direction), mutual information cost function, and spline final interpolation. To exclude areas outside the spinal canal which may move independently, the estimation of slice-wise motion parameters was restricted to a cylindric mask with a radius of 20 mm around the spinal cord centerline, created using the *sct_create_mask* function.

In the motion-corrected dataset, large intensity variations within the spinal cord between neighboring volumes were identified based on the DVARS metric (the spatial root-mean-square of the intensity difference image between successive volumes and within the spinal cord mask, as defined in (Power et al., 2012)) computed using FSL’s outlier detection tool (*fsl_motion_outliers*). Volumes with a DVARS exceeding the 75^th^ percentile + 1.5 times the interquartile range were considered outliers.

#### 2.5.2 Segmentation

After motion correction, the mean motion-corrected EPI image was manually segmented for spinal cord using FSL and was binarized with an inclusion threshold of 50%. The segmentation of the lower-resolution and lower-quality EPI image was guided by the segmentation of the higher-resolution and higher-quality ME-GRE image.

#### 2.5.3 Physiological noise modeling

Physiological noise regressors were obtained using the aCompCor method (Behzadi et al., 2007), implemented in the PhysIO toolbox (Kasper et al., 2017). The noise region of interest (ROI) was defined by manually segmenting the cerebrospinal fluid (CSF) in each slice. Special care was taken to exclude the interface between the CSF and the spinal cord from the segmentation to avoid partial volume effects, as well as the posterior part of the spinal canal due to the large number of nerves. No noise ROI was defined in the WM due to the risk of partial volume effects and signal mixing with the gray matter. For each slice, aCompCor extracted five principal components from the CSF ROI. The underlying assumptions for noise modeling in the CSF are that (i) CSF does not contain signal of interest (given the absence of neurons) and (ii) the physiological noise in the CSF shares common features with that in the gray matter.

#### 2.5.4 Co-registration to structural image

For each run, the mean motion-corrected EPI image was co-registered to the corresponding ME-GRE image using a non-linear slice-wise registration method in SCT (*sct_register_multimodal*). The co-registration was based on the spinal cord segmentation of both images, given the low contrast, and was conducted in two steps: first, using the centermass transformation (smooth factor of 3 mm, mean squares cost function), followed by the BSplineSyn transformation (Tustison & Avants, 2013) (10 iterations, mean squares cost function, and spline final interpolation). Note that the warping field was not applied to the EPI images directly; instead, it was applied to the parameter estimates during the subject-level analyses.

#### 2.5.5 Normalization to template

The spinal cord mask of the ME-GRE image was spatially normalized to the PAM50 spinal cord template using the *sct_register_to_template* function (De Leener et al., 2018) to obtain the forward (native to template space) warping field. We employed a two-label normalization approach, where we assigned labels to the slice with the largest spinal cord cross-sectional area within the lumbosacral enlargement (referred to as the “LSE landmark”) and the most caudal slice of the spinal cord (tip of the spinal cord) in the ME-GRE image. These labels were then matched to the corresponding spatial locations in the PAM50 template (disc labels 19 and 21) by the algorithm. The forward warping fields were used in the group-level analyses to warp statistical parametric maps from the native to the PAM50 space.

### 2.6 Image analysis and statistics

#### 2.6.1 Temporal signal-to-noise ratio

The temporal signal-to-noise ratio (tSNR) is a measure of the temporal stability of voxel time series within the EPI dataset. tSNR is commonly used to assess the quality of fMRI acquisitions as it is related to the minimal detectable effect size in the BOLD signal (Murphy et al., 2007). For each GE-EPI sequence, voxel-wise tSNR was computed on the processed EPI images (cropped and motion corrected, as described in Section 2.5.1), which further underwent linear detrending (i.e., first-order polynomial detrending) using the Matlab function *detrend*. To calculate tSNR, the mean of the voxel time series was divided by its standard deviation using the *sct_fmri_compute_tsnr* function in SCT. tSNR was only computed for the resting-state runs, as task runs may contain BOLD-related signal fluctuations. Subsequently, the tSNR values were averaged within the binary spinal cord mask, eroded by one voxel in the axial plane, to obtain a single tSNR value per slice. For slice-wise group statistics, axial slice stacks of individual subjects were aligned at their respective LSE landmark, i.e., the slice with the largest spinal cord cross-sectional area. Differences in tSNR across GE-EPI sequences and slices were assessed using two-way repeated measures ANOVA, with sequence and slice as within-subject variables. An interaction term between sequence and slice was not included, as we did not anticipate any influence of slice on sequence differences. Post-hoc tests with Tukey’s correction for multiple comparisons (p<0.05) were conducted to investigate pairwise differences.

#### 2.6.2 General linear model for task-based analysis

##### Run-level analysis

For each of the four GE-EPI sequences (Table 1), the processed EPI images of the task runs were spatially smoothed using an anisotropic 3D Gaussian kernel with a full-width-at-half-maximum of 1×1×5 mm^3^. In the run-level (first-level) analysis of these images, a general linear model (GLM) was fitted on each voxel time series using FSL fMRI Expert Analysis Tool (FEAT) (Woolrich et al., 2001). The GLM design matrix included three explanatory variables and several nuisance variables. The explanatory variables comprised a boxcar function describing the motion task (0 for rest, 1 for motion), which was convolved with three basis functions generated by FMRIB’s Linear Optimal Basis Sets (FLOBS) toolkit (Woolrich et al., 2004). Nuisance regressors included two motion parameters (x and y translational parameters output by *sct_fmri_moco*), motion outliers (output by *fsl_motion_outliers*), and the five principal components extracted from the CSF. The motion parameters and principal components served as slice-wise regressors (i.e., they varied from slice to slice) to account for variations in noise along the rostrocaudal direction. Besides model fitting, FSL FEAT also performed high-pass temporal filtering (cut-off frequency of 100 s) and pre-whitening (FSL FILM) on the voxel time series. As the rest block was not explicitly modeled, the contrast of parameter estimates (COPE) for the comparison (task > rest) corresponded to the parameter estimate of the first explanatory variable (β_1_).

##### Subject-level analysis

The two COPE maps from the run-level analyses (run 1 and 2) were warped to the ME-GRE image by applying the warping field obtained during co-registration (Section 2.5.4). The warped COPE maps were passed to the subject-level (second-level) analysis, which generated a subject-specific COPE map using a fixed effects model.

##### Group-level analysis

The subject-specific COPE maps were warped to the PAM50 space using the deformation field obtained during normalization (Section 2.5.5). Group-level (third-level) analysis was done by performing a non-parametric permutation test using FSL *randomise* with the default variance smoothing of 5 mm (Winkler et al., 2014). Notably, non-parametric tests generate the null distribution instead of modeling it, which results in exact inference (Eklund et al., 2016). In each permutation, the algorithm randomly flipped the signs of the subject-specific COPE values (n=12, resulting in 2^12^=4096 possible permutations), and performed a one-sample t-test on the sign-altered COPE values. On a voxel-by-voxel basis, the actual t-score was then tested against the distribution of the 4096 t-scores, yielding an (uncorrected) p-value. Threshold Free Cluster Enhancement (TFCE) was applied on the uncorrected maps of p-values to enhance cluster-like features in a threshold-free manner (without a cluster-defining threshold). The TFCE p-maps were family-wise error (FWE) corrected using three thresholds: (i) p_FWE_<0.05, following common thresholding practices, (ii) a more conservative threshold of p_FWE_<0.01 to emphasize differences in activation maps across the four sequences, and (iii) an even more conservative threshold of p_FWE_<0.001 to highlight only the voxels with the strongest association with the paradigm.

Three additional participants (1 female, 2 males, age (mean ± SD): 29.8 ± 3.1 years) were recruited as controls to evaluate the false positive rate of activation. These participants underwent the same scanning procedures but were instructed not to perform the task during the task runs. The control datasets were processed and analyzed in the same manner as described above.

#### 2.6.3 Distribution of BOLD activation

In the cross-section, the spinal cord was segmented into five ROIs: right ventral (RV), right dorsal (RD), left ventral (LV), left dorsal (LD), and medioventral (MV) spinal cord (Fig. 5A). The ROIs were created manually based on the PAM50 spinal cord atlas. The RV, RD, LV, and LD ROIs were delineated with a one-voxel gap between them. The MV ROI was created to separate the medial sulcal vein, a major vein draining the ventral gray matter horns, from the RV and LV ROIs. Along the rostrocaudal axis, the lumbosacral cord was segmented into approximate neurological levels using the relative lengths of neurological levels reported in a post-mortem study (Frostell et al., 2016) and assuming that the LSE landmark is at the border of L3 and L4 (Fig. 5B). To quantify BOLD activation, the mean t-score and the ratio of significant voxels (p_FWE_<0.01) were calculated within (i) each ROI over neurological levels L3-S2 and (ii) each neurological level over the whole spinal cord cross-section.

#### 2.6.4 BOLD effect size

The BOLD effect size was calculated as the percentage difference between the average signal intensities during motion and rest blocks within the task runs, for each participant and sequence. For each run, we generated the fully denoised dataset from the output of FSL FEAT. In particular, we summed up three 4D files: (i) the task-related signal, computed as the coefficient map associated with the first explanatory variable (task, β_1_, stored in the *pe1.nii.gz* file) multiplied with the HRF function (stored in the *design.mat* file), (ii) the offset signal intensity (stored in the *mean_func.nii.gz* file), (iii) and the residual errors (stored in the *res4d.nii.gz* file). Next, the mean time series was extracted within the RV ROI over the neurological levels L3 to S2, where motor-induced BOLD activation is expected from neuroanatomical considerations. For that, the RV ROI was transformed from the PAM50 to the native space using the backward warping fields obtained during co-registration and normalization. The mean time series were split into motion and rest blocks, where the first three data points of each block were omitted to exclude transient effects of the BOLD signal. Finally, we calculated the percentage difference between the mean signal intensities during motion and rest blocks.

## 3. Results

### 3.1 Image quality and temporal signal-to-noise ratio

Example slices of each GE-EPI sequence is provided in Fig. 3A. As expected, images acquired with a shorter TE display higher intensity levels. In addition, images with lower partial Fourier factor appear smoother (6/8 for OVS20 and iFOV28, 7/8 for iFOV35, 1 for iFOV42). In all participants, the OVS20 sequence produced folder-over artifacts at the anterior edge of the images despite the use of saturation bands (see green arrows in Fig 3A). Occasional signal dropouts at the dorsal edge of the spinal cord, induced by susceptibility artifacts, were more prominent at longer TE (see blue arrow in Fig. 3A).

**Fig. 3.**
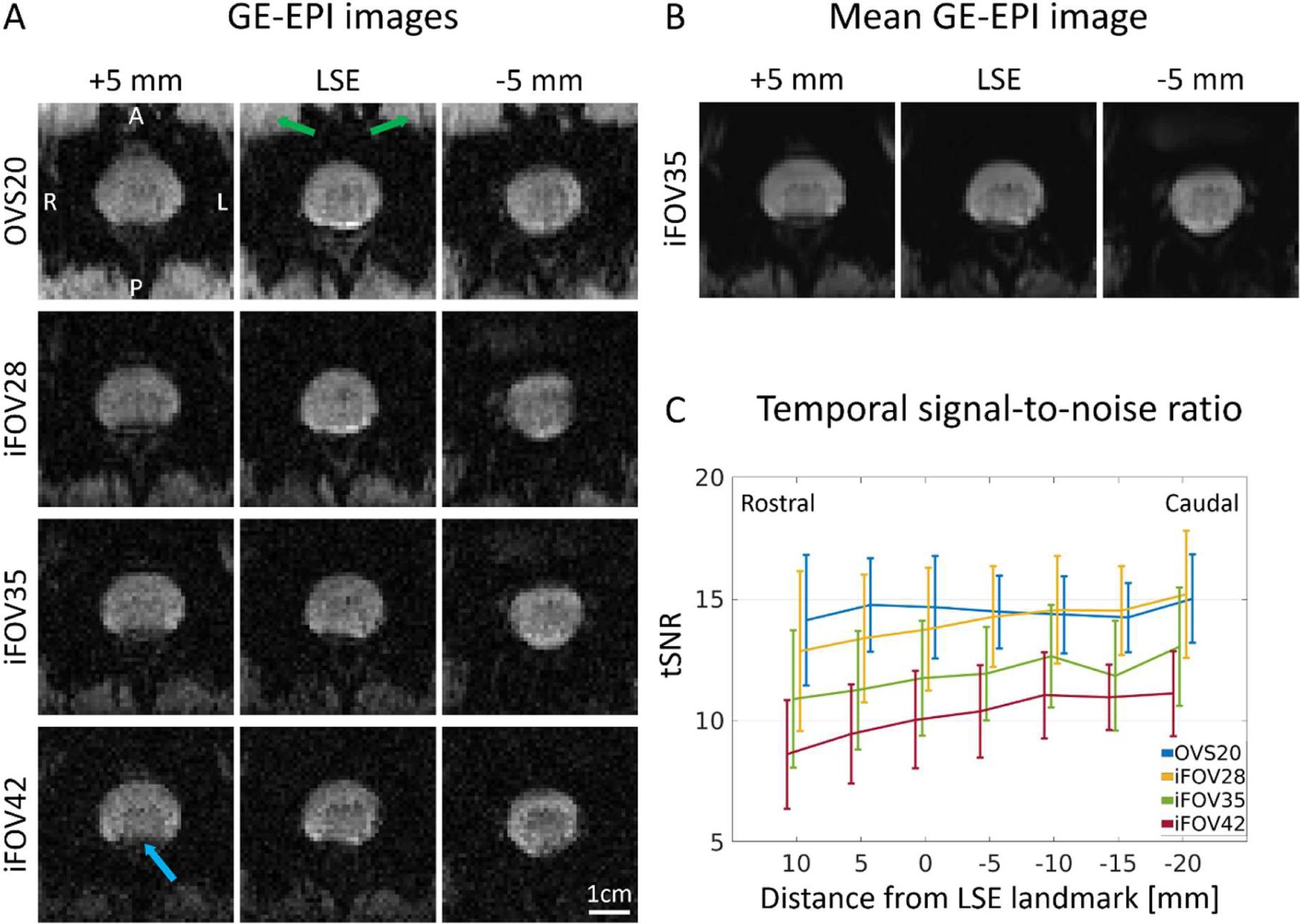
(A) Example slices from each GE-EPI sequence acquired in the same participant. For the OVS20 sequence, notice the fold-over artifact at the anterior edge of the image (indicated by the green arrows), resulting from imperfect saturation bands. Notice the higher smoothness of the images acquired with OVS20 and iFOV28 compared to those acquired with iFOV35 and iFOV42, which results from the application of 6/8 phase partial Fourier. The dorsal part of the spinal cord is occasionally affected by susceptibility artifacts (indicated by the blue arrow). (B) Mean image (averaged across all volumes) for the iFOV35 sequence. (C) Temporal signal-to-noise ratio (tSNR) within the spinal cord for each slice and GE-EPI sequence. Values represent group mean ± standard deviation (n=12). The individual axial slice stacks were aligned at the LSE landmark, i.e., the slice with the largest spinal cord cross-sectional area. A positive distance indicates a rostral direction from the LSE landmark. Displayed are slices where the full sample size (n=12) was available.

The tSNR values differed significantly between sequences (p<0.001) (Fig. 3C). Sequences with shorter TEs had higher tSNR, except for OVS20, which was not significantly different from iFOV28 (p=0.49). For example, at the LSE landmark, the tSNR was (group mean ± SD) 14.7 ± 2.1 for the OVS20, 13.8 ± 2.5 for the iFOV28, 11.8 ± 2.4 for the iFOV35, and 10.0 ± 2.0 for the iFOV42 sequence. Slices also had a significant effect on tSNR (p=0.007), showing an increasing tendency in the caudal direction (Fig. 3C).

On average, the number of outlier volumes ranged from 10.9 to 20.2 per run, although with large inter-subject variability (Table S1). For each sequence, the second run had, on average, 2-3 more outlier volumes than the first run.

### 3.2 Distribution of BOLD activation

#### Group-level results

Group-level parametric maps are presented in Fig. 4A and B, while plots illustrating the ratio of significant (supra-threshold) voxels and mean t-scores across ROIs and neurological levels are displayed in Fig. 4C and Fig. 5. Numerical values are provided in Table 2. For each GE-EPI sequence, a cluster of activation was found in the right (ipsilateral) ventral region of the spinal cord, which also extended into the right dorsal region (Fig. 4A). This was also evident in the high ratio of significant voxels (p_FWE_<0.01) and high t-score in the right ventral and right dorsal ROIs (Fig. 5A). At the same threshold, little to no activation was observed in the left (contralateral) ventral and dorsal ROIs. Along the rostrocaudal axis, significant voxels were detected between L3-S2, with the highest ratio of significant voxels and highest mean t-score at L5 (Fig. 4C, Fig. 5B).

**Fig 4.**
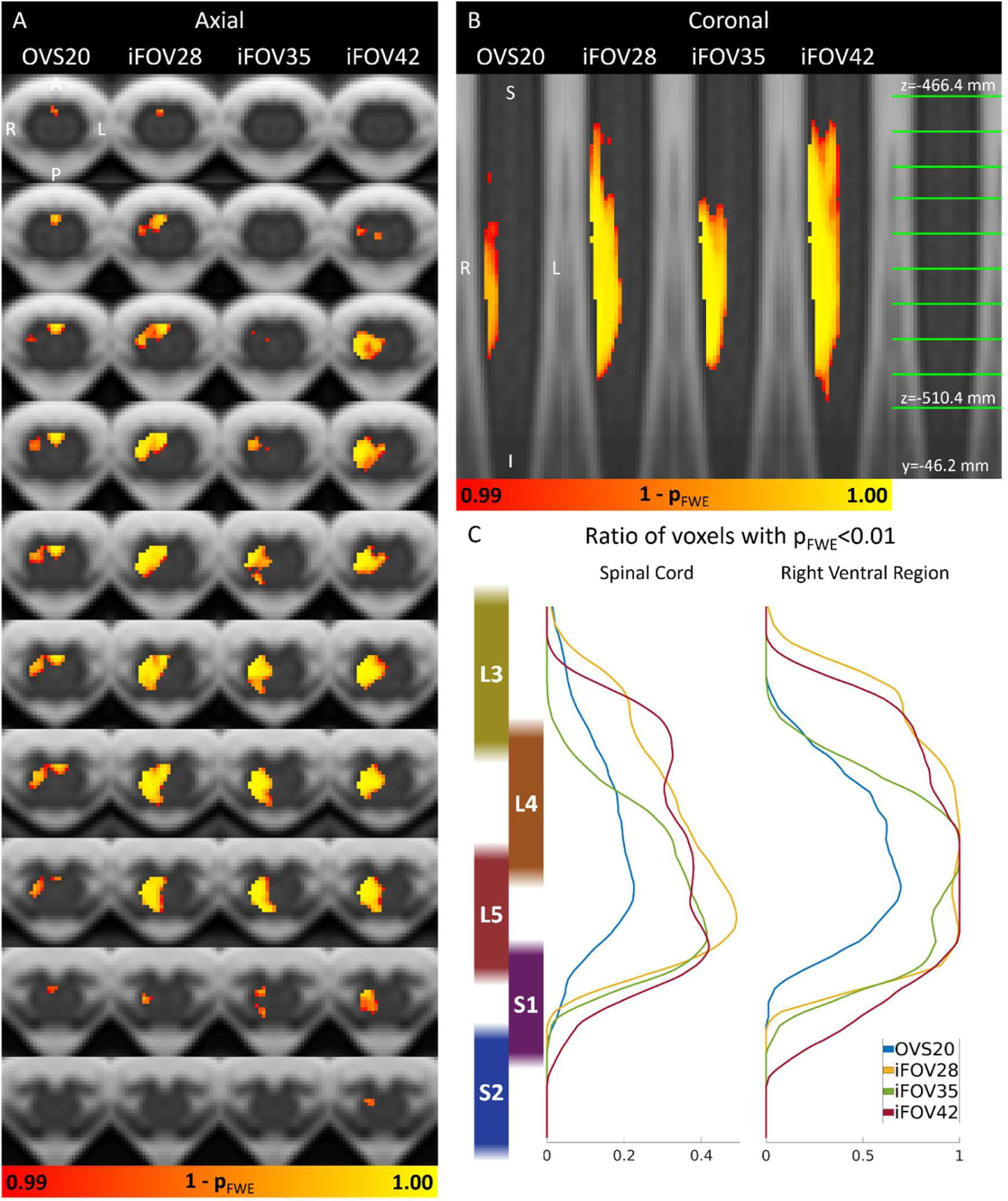
Group-level statistical parametric maps (n=12) representing the family-wise error corrected p-value (p_FWE_) for the contrast task vs. baseline. Parametric maps are thresholded at p_FWE_ < 0.01 and shown in both the axial (A) and coronal (B) planes as heatmaps superimposed on the PAM50 template. The axial slices are evenly spaced between z=-466.4 mm (most rostral slice, PAM50 coordinates) and z=-510.4 mm (indicated by green lines in the coronal plane). Images are displayed in radiological convention. (C) Rostrocaudal distribution of significant voxels (p_FWE_ < 0.01) within the whole spinal cord and the ipsilateral (right) ventral region of the spinal cord (see Fig. 5A for definition). Spinal cord neurological levels are approximated using the relative proportions of neurological levels reported in (Frostell et al., 2016).

**Fig. 5.**
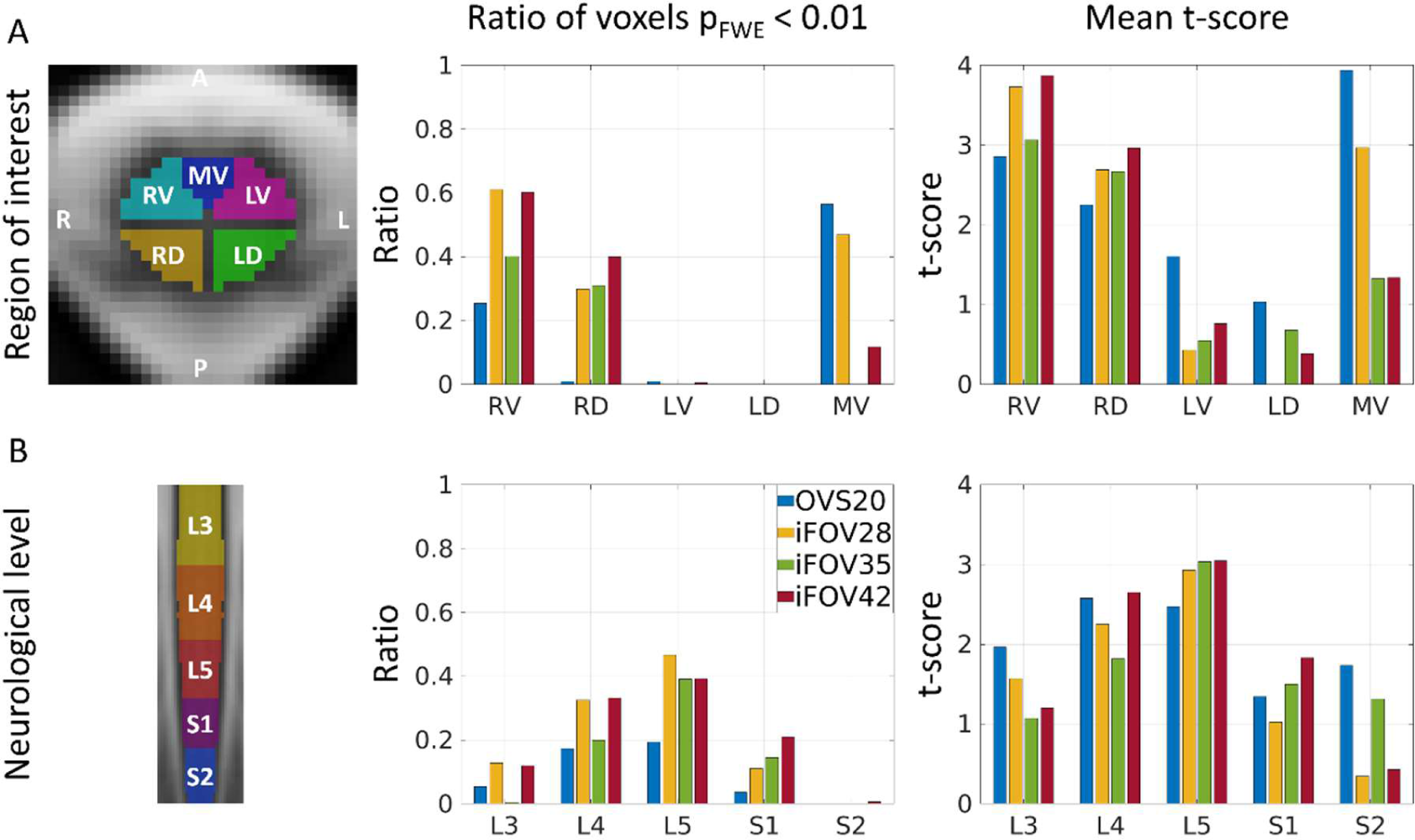
Ratio of significant voxels (p_FWE_ < 0.01) and mean t-score within (A) each region of interest over neurological levels L3-S2 and (B) each neurological level over the whole spinal cord cross-section. The regions of interest are shown for a single slice and included the right ventral (RV), right dorsal (RD), left ventral (LV), left dorsal (LD), and medioventral (MV) spinal cord. Spinal cord neurological levels are approximated using the relative proportions of neurological levels reported in (Frostell et al., 2016).

**Table 2.** Mean t-score and ratio of voxels with p_FWE_ < 0.05, 0.01, and 0.001 within each region of interest (over neurological levels L3-S2) and each neurological level (over the whole spinal cord cross-section).

|  | Sequence | Region of interest |  |  |  |  | Neurological level |  |  |  |  |
| --- | --- | --- | --- | --- | --- | --- | --- | --- | --- | --- | --- |
|  |  | RV | RD | LV | LD | MV | L3 | L4 | L5 | S1 | S2 |
| t-score<br>(mean) | OVS20 | 2.85 | 2.25 | 1.60 | 1.03 | 3.94 | 1.97 | 2.58 | 2.47 | 1.34 | 1.73 |
|  | iFOV28 | 3.73 | 2.69 | 0.43 | 0.01 | 2.97 | 1.57 | 2.25 | 2.93 | 1.02 | 0.35 |
|  | iFOV35 | 3.07 | 2.67 | 0.55 | 0.68 | 1.33 | 1.07 | 1.82 | 3.04 | 1.50 | 1.31 |
|  | iFOV42 | 3.87 | 2.96 | 0.76 | 0.38 | 1.34 | 1.20 | 2.65 | 3.05 | 1.83 | 0.43 |
| $p_{FWE} < 0.05$<br>(ratio) | OVS20 | 0.58 | 0.33 | 0.10 | 0.02 | 0.84 | 0.22 | 0.45 | 0.42 | 0.20 | 0.01 |
|  | iFOV28 | 0.81 | 0.56 | 0.01 | 0.00 | 0.61 | 0.30 | 0.44 | 0.53 | 0.26 | 0.01 |
|  | iFOV35 | 0.58 | 0.47 | 0.00 | 0.01 | 0.08 | 0.07 | 0.31 | 0.49 | 0.25 | 0.01 |
|  | iFOV42 | 0.77 | 0.63 | 0.04 | 0.00 | 0.28 | 0.24 | 0.45 | 0.51 | 0.34 | 0.03 |
| $p_{FWE} < 0.01$<br>(ratio) | OVS20 | 0.25 | 0.01 | 0.01 | 0.00 | 0.57 | 0.05 | 0.17 | 0.19 | 0.04 | 0.00 |
|  | iFOV28 | 0.61 | 0.30 | 0.00 | 0.00 | 0.47 | 0.13 | 0.33 | 0.47 | 0.11 | 0.00 |
|  | iFOV35 | 0.40 | 0.31 | 0.00 | 0.00 | 0.00 | 0.00 | 0.20 | 0.39 | 0.14 | 0.00 |
|  | iFOV42 | 0.60 | 0.40 | 0.01 | 0.00 | 0.12 | 0.12 | 0.33 | 0.39 | 0.21 | 0.01 |
| $p_{FWE} < 0.001$<br>(ratio) | OVS20 | 0.00 | 0.00 | 0.00 | 0.00 | 0.13 | 0.01 | 0.03 | 0.01 | 0.00 | 0.00 |
|  | iFOV28 | 0.23 | 0.07 | 0.00 | 0.00 | 0.18 | 0.01 | 0.14 | 0.21 | 0.01 | 0.00 |
|  | iFOV35 | 0.16 | 0.10 | 0.00 | 0.00 | 0.00 | 0.00 | 0.04 | 0.22 | 0.03 | 0.00 |
|  | iFOV42 | 0.27 | 0.10 | 0.00 | 0.00 | 0.00 | 0.01 | 0.13 | 0.21 | 0.03 | 0.00 |

FWE, family-wise error corrected; LD, left dorsal; LV, left ventral; MV, medioventral; RD, right dorsal; RV, right ventral.

We observed differences in the level and extent across the GE-EPI sequences. The ratio of significant voxels (p_FWE_<0.01) and the mean t-score in both the right ventral and dorsal ROIs tended to increase with longer TE (Fig. 5A). Notably, the OVS20 sequence showed only a minimal number of significant voxels in the right dorsal ROI. Sequences with shorter TE (20 and 28 ms) displayed a cluster of activation in the medioventral ROI, which was not present at longer TE (35 and 42 ms) (Fig. 4A, Fig. 5A).

#### Subject-level results

Subject-level parametric maps were noisier but displayed clusters of significant voxels in all participants (Fig. S2). Participants undergoing the task-free paradigm exhibited a minimal and spatially random pattern of activation (Fig. S2), with false positive rates of 1.40 ± 0.75% for the OVS20, 1.10 ± 0.26% for the iFOV28, 1.20 ± 0.14% for the iFOV35, and 1.15 ± 0.08% for the iFOV42 sequence when applying a threshold of z > 2.3263 (p < 0.01).

### 3.3 BOLD effect size

The BOLD effect size showed an increasing trend with increasing TE, with values (group mean ± SD) of 0.34 ± 0.21% for the OVS20, 0.35 ± 0.19% for the iFOV28, 0.42 ± 0.26% for the iFOV35, and 0.58 ± 0.27% for the iFOV42 sequence.

## Discussion

Task-related BOLD activation associated with ankle movement is detected in the locations expected from neuroanatomical considerations, which demonstrates the feasibility and validity of lumbosacral BOLD fMRI by using reduced-FOV single-shot gradient-echo echo planar imaging (GE-EPI) sequences. While all investigated GE-EPI sequences detected neuronal activation, the one with the longest echo time (TE=42 ms) yielded the highest BOLD effect size. Sequences with TE below 30 ms were found to be susceptible to spatially unspecific BOLD activation due to large vein effects. The overall small BOLD effect size of approximately 0.5% underscores the necessity for effective denoising. Further research aimed at optimizing spinal cord fMRI has the potential to improve the sensitivity and robustness of lumbosacral fMRI.

### 3.4 Lateralized BOLD activation in response to unilateral lower extremity motor task

The primary cluster of BOLD activation, induced by unilateral (right-sided) ankle plantar and dorsiflexion, was extremely lateralized on the ipsilateral (right) side, aligning with our current understanding of spinal cord organization. Minimal to no activation was detected on the contralateral (left) side. Lateralization was also observed in previous fMRI studies involving upper extremity motor tasks, such as wrist flexion (Weber et al., 2016b), wrist extension, wrist abduction, and finger abduction (Kinany et al., 2019).

In the cross-section, the BOLD activation was centered in the ipsilateral ventral part of the spinal cord. The spatial extent of this cluster depended on the applied threshold. At a conservative threshold (p_FWE_< 0.001), the cluster was mainly localized within the ipsilateral ventral gray matter only (Fig. S1B), while extending further into the ipsilateral dorsal region at more liberal thresholds (p_FWE_<0.01, Fig. 4, and p_FWE_<0.05, Fig. S1A). This spatial distribution aligns with neuroanatomy, as lower motor neurons are primarily located in the ventral gray matter horn (McHanwell & Watson, 2009). Moreover, ankle movements also activate the interneurons of the proprioceptive pathways, which may contribute to the observed (weak) activation in the dorsal horns (Marmigère & Ernfors, 2007).

BOLD activation is not confined to the gray matter but extends into the surrounding white matter, as particularly evident at a liberal threshold (p_FWE_<0.05, Fig. S1A). On one hand, the BOLD contrast arises from the venous capillaries and other draining vessels surrounding the activated neurons (Keilholz et al., 2006; Kennerley et al., 2010), limiting the spatial specificity and representing an inherent limitation of the technique. On the other hand, smoothing the images introduces further blurring of the activation patterns.

The motor task involved ankle dorsiflexion and plantar flexion, which are executed by myotomes L4 and S1, respectively. Indeed, all sequences detected activation ranging from L3 to S2, with a single peak observed at L5. We argue that the absence of two distinct peaks, reflecting ankle dorsiflexion and plantar flexion, can be attributed to the inter-subject variation in nerve root muscle innervation along with the limited rostrocaudal resolution (5 mm slice thickness), which blurs the two peaks into a single one at group-level. Furthermore, studies have demonstrated that the involved muscles can be activated from various nerve roots (Hashimoto et al., 2023; Lorach et al., 2023; Phillips & Park, 1991).

### 3.5 Group activation maps are influenced by the echo time

Although BOLD activations were detected in the ipsilateral ventral gray matter horn (the expected location) using each of the four GE-EPI sequences (with TE of 20, 28, 35, and 42 ms), differences in the level and spatial pattern of these activations were also observed. At TE below 30 ms (sequences OVS20 and iFOV28), a focal activation was identified in the medioventral region of the spinal cord. Notably, at the shortest investigated TE (20 ms), this cluster exhibited even stronger activation than the one in the ipsilateral ventral region and was the only one to survive a more conservative threshold of p_FWE_<0.001. We argue that this cluster results from task-induced susceptibility changes in large draining vessels, indirectly reflecting neuronal activation in the ventral gray matter horns. First, the location of the cluster corresponds to that of the medial sulcal vein, the major venous vessel draining the ventral gray matter horns (Thron, 2016). Second, task-free controls did not exhibit similar activations. In fact, previous studies have highlighted the role of large draining vessels in contributing to the BOLD contrast (Engel et al., 1997; Lai et al., 1993; Uludaǧ et al., 2009). Consistent with our findings, the contribution of large veins to the BOLD contrast has been reported to be higher at shorter TE values (Triantafyllou et al., 2011).

We also observed an increase in the BOLD effect size within the ipsilateral ventral region with longer echo times, ranging from 0.35% (TE=20 ms) to 0.58% (TE=42 ms). Similarly, the ratio of significant voxels and the mean t-score within the same region tended to increase with longer echo times. We note, however, that these latter changes were not monotonic, as iFOV35 yielded lower values that iFOV28. We argue that the values obtained with a TE of 28 ms are inflated due to the strong activation from the neighboring medioventral region (large vein effect). We also note that the BOLD effect sizes reported here are lower than those reported in the cervical cord using GE-EPI at 3T (Barry et al., 2021; Weber et al., 2020); however, the specific values are highly sensitive to the methodology employed for their computation.

In theory, the BOLD effect size in a given voxel exhibits a broad peak, reaching its maximum when the TE approximately corresponds to the transversal relaxation time (T_2_*) within that voxel. Although this relationship often does not hold in practice due to the presence of thermal and physiological artifacts (Fera et al., 2004), the broad peak ensures that BOLD contrast is detectable within a relatively wider range of TE values around the optimal value (Menon et al., 1993; van der Zwaag et al., 2009). While we are not aware of any reports on T_2_* values in the lumbosacral cord, T_2_* was measured to be 41.3 ± 5.6 ms in the gray matter of the cervical spinal cord at 3T (Barry & Smith, 2019). This measurement aligns with our observation of the highest BOLD effect size occurring at a TE of 42 ms. It is worth noting that T_2_* values measured in the cervical cord are shorter compared to those in the brain cortex (Barry & Smith, 2019), emphasizing that a TE optimized for the brain might not be optimal for the spinal cord.

Establishing a single optimal TE is challenging in practice, as the local T_2_* depends on various factors, including shimming quality and sequence parameters such as voxel size. Moreover, there is a significant spatial variability in T_2_* both in the brain and spinal cord due to differences in magnetic susceptibility among surrounding tissues (Barry & Smith, 2019). Nevertheless, with second-order shimming and a resolution of 1×1×5 mm^3^, our findings suggest an optimum TE of approximately 42 ms for detecting spatially specific BOLD activation within the ipsilateral (right) ventral gray matter. This TE is longer than those typically used in recent spinal cord BOLD fMRI studies. We note that a longer TE might detect even higher BOLD activation. However, an increase in TE also prolongs the repetition time (TR), resulting in a reduction in the number of acquired volumes within the same imaging time, which, in turn, reduces the power of the fMRI experiment (Murphy et al., 2007).

### 3.6 Considerations for image acquisition

Several early spinal cord fMRI studies utilized the SEEP contrast to detect task-related neuronal activation (Ghazni et al., 2010; Kornelsen & Stroman, 2004; Ng et al., 2006); however, the SEEP contrast has been the subject of debate (Jochimsen et al., 2005). Additionally, the GE-EPI sequence, which targets the BOLD contrast, demonstrated higher sensitivity, spatial specificity, and reproducibility compared to the turbo spin-echo fMRI sequence, which targets the SEEP contrast (Bouwman et al., 2008; Jochimsen et al., 2005).

The reduced-FOV single-shot GE-EPI sequences utilized in this study were adapted from sequences employed in previous studies (Kinany et al., 2019, 2022; Weber et al., 2016a, 2016b, 2020), where the sequence’s sensitivity to detect BOLD activation in the spinal cord was established. The in-plane resolution and matrix size were also comparable to the spinal cord diffusion MRI sequence in the cervical MRI consensus protocol (Cohen-Adad et al., 2021a).

We argue that tSNR serves as a valuable tool for evaluating the quality of shimming and image processing steps, and for estimating the statistical power of the fMRI experiment (Murphy et al., 2007). However, the observation that the sequence with the lowest tSNR (iFOV42) yielded the higher BOLD effect size clearly demonstrates that tSNR alone is inadequate for assessing the sequences’ sensitivity to BOLD signal. For example, caution is necessary when comparing sequences with different partial Fourier acceleration factors. The application of partial Fourier introduces smoothing in the image, which increases the *apparent* tSNR but reduces the intrinsic tSNR. Therefore, the observed increases in tSNR when using shorter TE are attributed not only to T_2_* decay but also to the application of partial Fourier. Note that for the iFOV sequences, the shorter TE also lead to a shorter TR (with adjusted flip angle), which counteracts the tSNR increase; however, this effect appeared to be minor. Despite the shorter TE and longer TR, the OVS20 sequence yielded similar results to iFOV28, suggesting that, given the same TE and TR, inner field-of-view excitation would result in higher tSNR. This may be due to the imperfect saturation band, causing aliasing in the ventral part of the image (Fig. 3A), which might generate ghosts overlaying the spinal cord.

While not statistically significant, sequences with longer TE tended to show an increased number of outlier volumes. Considering the counterbalanced order of sequences, this trend might not be attributed to a higher level of motion toward the end of the imaging session but rather to the lower tSNR and higher artifact level associated with longer TE. Furthermore, the smoother appearance of the images when applying partial Fourier might reduce image variation and hence the number of detected outliers.

### 3.7 Limitations and future directions

Automatic segmentation techniques for the spinal cord in EPI-based fMRI data are yet to be developed and validated, leaving manual segmentation as the current standard procedure. This is a recognized issue within the research community and an ongoing joint effort is underway (Bautin & Cohen-Adad, 2021). While manual segmentation leads to intra- and inter-rater variability in the processing pipeline, it has been demonstrated that this variability does not introduce systematic bias into the group-level fMRI activation maps (Hoggarth et al., 2022).

A challenge specific to lumbosacral cord imaging is the difficulty of identifying neurological levels in vivo, given the mismatch between vertebral and neurological levels in the lumbosacral cord (Canbay et al., 2014), which precludes the use of vertebral levels as neuroanatomical landmarks. While the two-label normalization approach (Section 2.5.5) ensured the alignment of the center of the lumbosacral enlargement and the tip of the spinal cord across participants, this approach does not ensure the alignment of neurological levels in between these two landmarks. For quantifying BOLD-activation at specific neurological levels, we divided the space between the LSE landmark and the tip of the spinal cord according to the relative lengths of neurological levels reported in a postmortem study (Frostell et al., 2016). However, even if we could identify neurological levels, one must be aware of the considerable between-subject variation in the distribution of the targeted neuronal populations across neurological levels. For example, the rostrocaudal location of two muscle projectomes were even found to be inverted in an individual with spinal cord injury (Rowald et al., 2022).

We used the canonical HRF implemented in FSL, which was originally devised for the brain, for modeling the BOLD-related signal changes in the general linear model. However, the HRF might be different between the brain and in the spinal cord but even among different parts of the spinal cord (Giulietti et al., 2008), compromising the use of a single HRF across the entire FOV. While we addressed variations from the canonical HRF by using FLOBS in the general linear model, residual differences might still affect the group-level analyses (Handwerker et al., 2004).

Although visual cues were given to participants to initiate movements, we did not measure the muscle forces associated with ankle dorsi- and plantar flexion due to the unavailability of MRI-compatible dynamometers for lower extremity tasks. Therefore, drawing associations between muscle force and BOLD activity was not feasible.

In this study, we compared sequences with varying echo times in terms of their ability to detect task-related activity. Comparison of these sequences in terms of their ability to detect resting-state functional connectivity will be the focus of future investigations. Notably, a previous study performed in the cervical cord has observed differences in dynamic functional connectivity patterns when employing different sequence parameters (Kinany et al., 2022).

Finally, it is important to note the considerable inter-subject variability in both the level and extent of BOLD activation (Fig. S2). We observed that subjects with a higher number of outlier volumes tended to have fewer significant voxels (Fig. S3). Interestingly, individuals who exhibited many outliers were mostly those with the least MRI experience. This suggests that those with more MRI experience might find it easier to relax and avoid motion artifacts. Nevertheless, additional sources of inter-subject variability remain unknown and warrant further investigation. Moreover, a more in-depth examination of lumbosacral fMRI within a test-retest setting is necessary.

### 3.8 Conclusions

We demonstrated the feasibility of detecting motor-evoked neuronal activation within the lumbosacral cord using BOLD fMRI in combination with a block-design lower extremity motor task. The detected activation is in line with the expected location from neuroanatomical considerations. We further highlight the importance of echo time in the sensitivity to BOLD changes in the lumbosacral cord. Functional MRI in the lumbosacral cord has significant implications for assessing motor function and its alterations in disease or after spinal cord injury.

## Author Contributions

C.W.K.: Conceptualization, Data Acquisition, Methodology, Formal Analysis, Visualization, and Writing (Original draft, Review & Editing).

J.F.: Methodology and Writing (Review & Editing).

P.F.: Conceptualization, Funding Acquisition, Supervision, and Writing (Review & Editing).

G.D.: Conceptualization, Data Acquisition, Methodology, Formal Analysis, Funding Acquisition, Supervision, and Writing (Original draft, Review & Editing).

## Funding

This work was funded by the Swiss National Science Foundation (SNSF) (32003B_204934) and the Swiss Paraplegic Foundation (Foko_2020_03). PF is funded by a SNSF Eccellenza Professorial Fellowship Grant (PCEFP3_181362/1).

## Declaration of Competing Interests

The authors do not have any competing interests to declare.

## Acknowledgements

We thank all the volunteers for participating in this study. Imaging was performed with support of the Swiss Center for Musculoskeletal Imaging, SCMI, Balgrist Campus AG, Zürich.

## Supplementary material

**Fig. S1.**
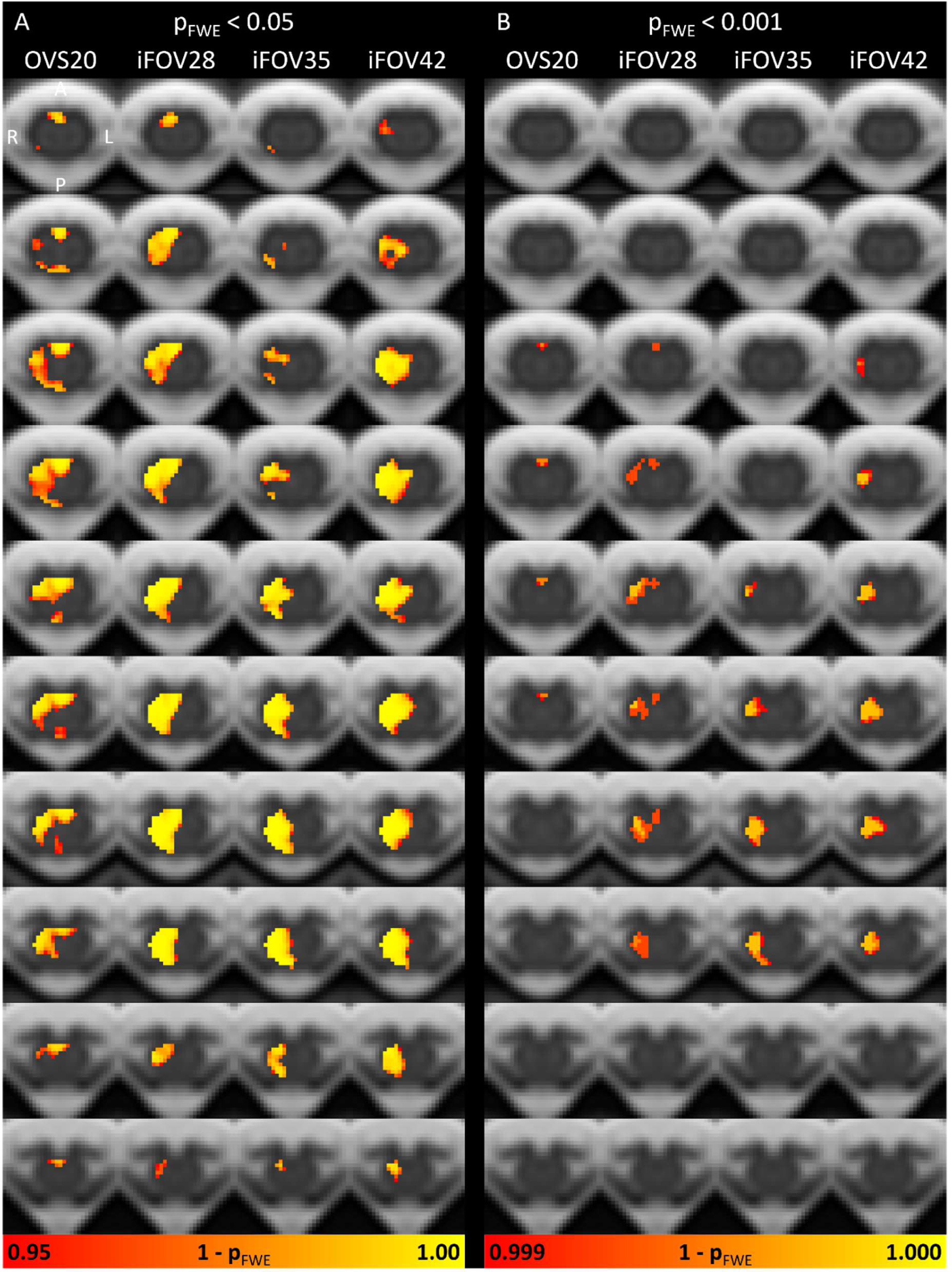
Group-level statistical parametric maps (n=12) representing the family-wise error corrected p-value (p_FWE_) for the contrast task vs. baseline. Parametric maps are thresholded at (A) p_FWE_ < 0.05 and (B) p_FWE_ < 0.001 and shown in the axial plane as heatmaps overlaid on the PAM50 template for each GE-EPI sequence. The axial slices are evenly spaced between z=-466.4 mm (most rostral slice, PAM50 coordinates) and z=-510.4 mm. Images are displayed in radiological convention.

**Fig. S2.**
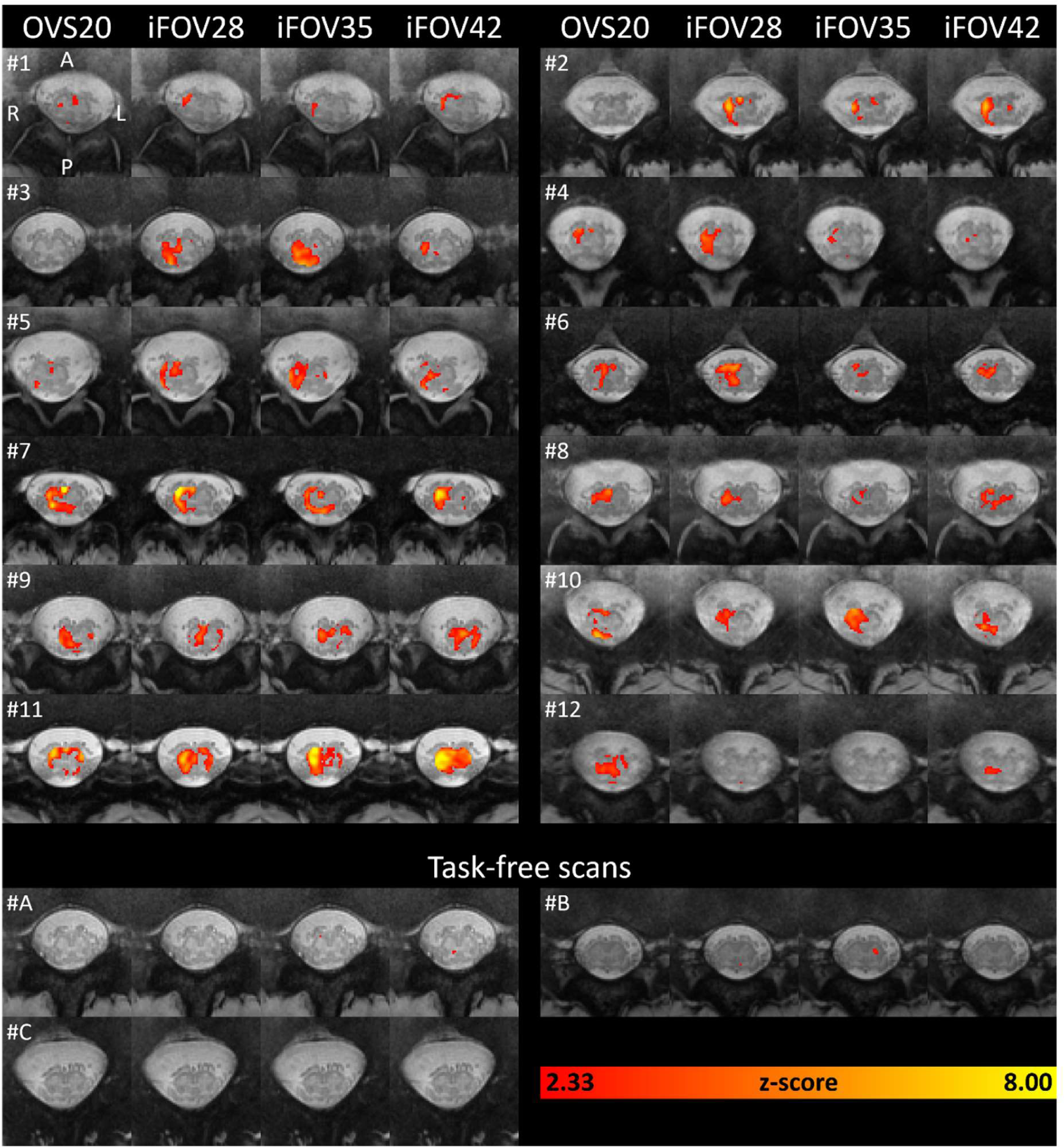
Subject-level statistical parametric maps representing the z-score for the contrast task vs. baseline. Parametric maps are thresholded at z > 2.33 (corresponding to uncorrected p < 0.01) and shown in the axial plane as heatmaps overlaid on the PAM50 template for each GE-EPI sequence. Three subjects underwent task-free scans (i.e., the same paradigm but without performing the task). For each subject, the slice located 10 mm (2 slices) caudal to the slice with the largest spinal cord cross-sectional area is shown. Images are displayed in radiological convention.

**Fig. S3.**
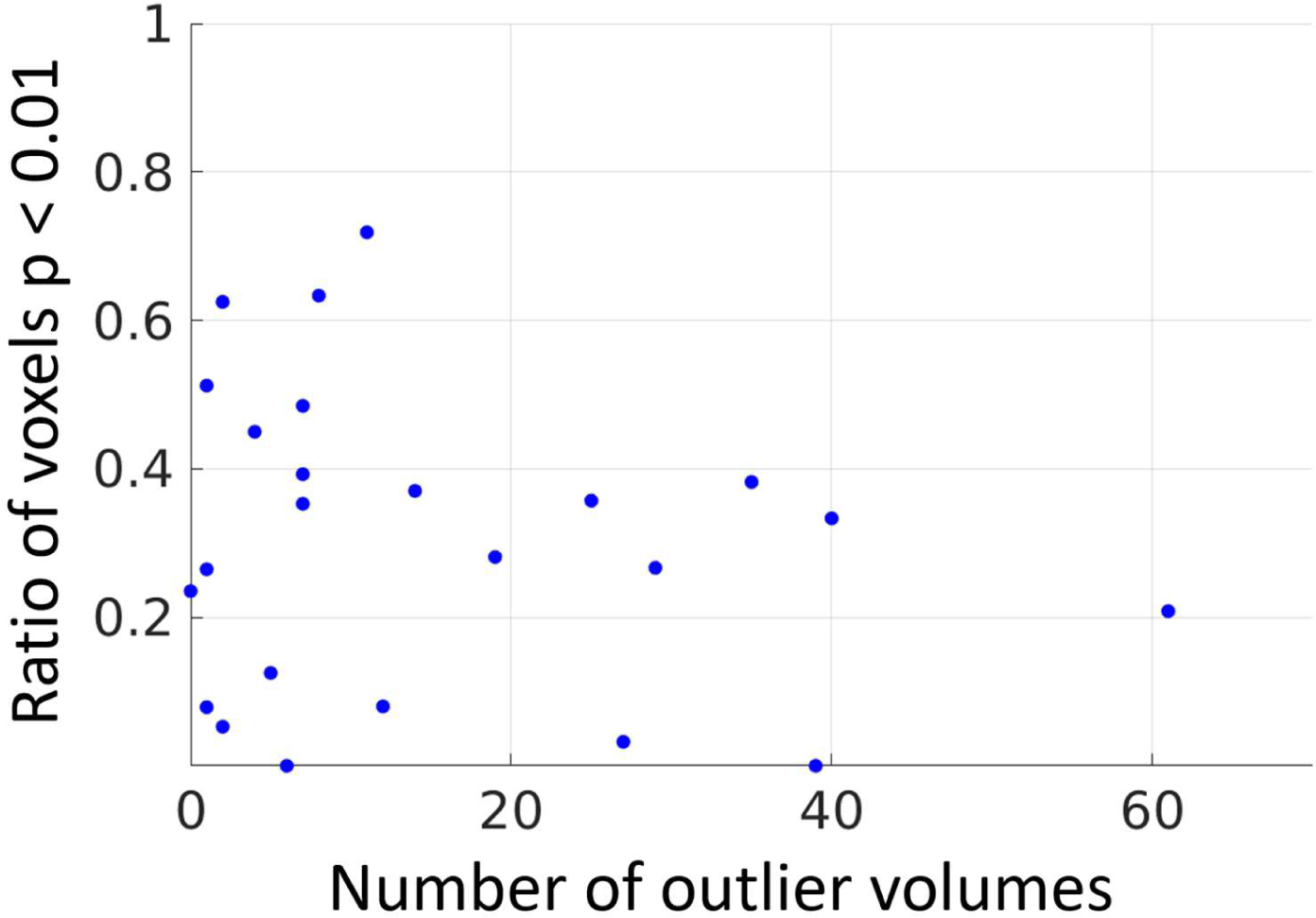
Scatter plot illustrating the correlation between the ratio of the significant voxels within the ipsilateral (right) ventral region (see Fig. 5A for definition) and the number of outlier volumes detected by *fsl_motion_outliers*. Each data point represents an individual subject. Significant voxels were defined as those with a subject-level z-score above 2.33 (corresponding to an uncorrected p < 0.01). For each subject, the number of outlier volumes was averaged across both runs of the iFOV42 sequence (n=429 volumes in each run). While a low number of outliers (i.e., below 20) did not appear to be associated with the ratio of significant voxels, a high number of outliers tended to correspond to fewer significant voxels. While not shown here, sequences OVS20, iFOV28, and iFOV35 displayed a qualitatively similar pattern.

**Table S1.** Number of outliers detected by *fsl_motion_outliers* for each run.

| Subject | OVS20 (n=411) |  | iFOV28 (n=505) |  | iFOV35 (n=466) |  | iFOV42 (n=429) |  |
| --- | --- | --- | --- | --- | --- | --- | --- | --- |
|  | Run 1 | Run 2 | Run 1 | Run 2 | Run 1 | Run 2 | Run 1 | Run 2 |
| S01 | 44 | 38 | 20 | 44 | 31 | 46 | 27 | 39 |
| S02 | 16 | 28 | 25 | 8 | 25 | 26 | 35 | 25 |
| S03 | 6 | 12 | 14 | 8 | 28 | 47 | 7 | 7 |
| S04 | 6 | 7 | 6 | 5 | 11 | 5 | 4 | 12 |
| S05 | 8 | 3 | 2 | 2 | 3 | 6 | 0 | 1 |
| S06 | 8 | 36 | 12 | 17 | 15 | 13 | 14 | 7 |
| S07 | 5 | 3 | 1 | 2 | 0 | 6 | 1 | 2 |
| S08 | 14 | 13 | 7 | 14 | 17 | 22 | 1 | 2 |
| S09 | 20 | 15 | 16 | 24 | 19 | 15 | 19 | 29 |
| S10 | 6 | 15 | 6 | 11 | 32 | 29 | 40 | 61 |
| S11 | 7 | 3 | 6 | 6 | 6 | 8 | 8 | 11 |
| S12 | 3 | 2 | 16 | 14 | 19 | 19 | 5 | 6 |
| Mean | 11.9 | 14.6 | 10.9 | 12.9 | 17.2 | 20.2 | 13.4 | 16.8 |
| SD | 11.3 | 12.8 | 7.4 | 11.7 | 10.7 | 14.7 | 13.8 | 18.4 |

## Notes

### Competing Interest Statement

The authors have declared no competing interest.

